# Understanding hydropower impacts on Amazonian wildlife is limited by a lack of robust evidence: results from a systematic review

**DOI:** 10.1101/2021.07.01.450737

**Authors:** Eduardo Rodrigues dos Santos, Fernanda Michalski, Darren Norris

**Author notes:** Corresponding Author: Darren Norris, Ecology and Conservation of Amazonian Vertebrates Research Group, Federal University of Amapá, Rod. Juscelino Kubitschek Km 02, 68903-419 Macapá, Brazil.

## Abstract

**Background and Research Aims:** Although hydropower provides energy to fuel economic development across Amazonia, strategies to minimize or mitigate impacts in highly biodiverse Amazonian environments remain unclear. The growing number of operational and planned hydroelectrics requires robust scientific evidence to evaluate impacts of these projects on Amazonian vertebrates. Here we investigated the existing scientific knowledge base documenting impacts of hydropower developments on vertebrates across Brazilian Amazonia.

**Methods:** We reviewed the scientific literature from 1945 to 2020 published in English, Spanish and Portuguese to assess the temporal and spatial patterns in publications and the types of study design adopted as well as scientific evidence presented.

**Results:** A total of 24 published articles documented impacts on fish (n = 20), mammals (n = 3) and freshwater turtles (n = 1). Most study designs (87.5%) lacked appropriate controls and only three studies adopted more robust Before-After-Control-Impact designs. The published evidence did not generally support causal inference with only two studies (8.3%) including appropriate controls and/or confounding variables.

**Conclusion:** Decades of published assessments (54.2% of which were funded by hydropower developers or their subsidiaries) do not appear to have established robust evidence of impacts of hydropower developments on Amazonian vertebrates. This lack of robust evidence could limit the development of effective minimization and mitigation actions for the diverse vertebrate groups impacted by hydroelectrics across Brazilian Amazonia.

**Implications for Conservation:** To avoid misleading inferences there is a need to integrate more robust study designs into impact assessments of hydropower developments in the Brazilian Amazon.

## Introduction

The development and operation of hydroelectric power plants generates multiple environmental and social impacts across tropical regions, ranging from habitat destruction to changes in river flow, habitat fragmentation, and overhunting (Aurelio-Silva et al., 2016; Benchimol & Peres, 2015; Bueno & Peres, 2019; Cosson et al., 1999; Palmeirim et al., 2017). The increasing number of hydroelectrics in tropical rivers means there is an urgent need to understand impacts to establish minimization and mitigation actions necessary to ensure sustainability of these developments. To date evidence documenting impacts is limited, for example the only synthesis at the Environmental Evidence database is on impacts to fish mortality (Algera et al., 2020) and fish productivity (Rytwinski et al., 2020) in temperate regions (https://environmentalevidence.org/completed-reviews/?search=dam, accessed 14 July 2021).

In South America, hydropower projects with reservoirs and run-of-river dams are common (Finer & Jenkins, 2012). For example, in 2021 Brazilian Amazonia has 29 operational hydroelectric power plants (including only those with installed power > 30 MW) and an additional 93 in process of regularization and construction (SIGEL, 2021). Projects with reservoir storage (e.g. Balbina dam in Brazil), make it possible to adjust the level of water to produce energy during periods of water scarcity, which can make substantial changes to both the landscape and water flow (Egré & Milewski, 2002; Fearnside, 1989). Projects using run-of-river dams use the natural river flow to generate energy and can therefore reduce environmental impacts in certain cases (Egré & Milewski, 2002). Yet due to highly seasonal rainfall and river flow rates the vast majority of Amazonian run-of-river dams include reservoirs e.g. Belo Monte (Fearnside, 2006; Hall & Branford, 2012) and as such generate drastic impacts on flowrates (Mendes et al., 2021).

The Amazon rainforest is renowned for its globally important biodiversity and availability of hydric resources (Dirzo & Raven, 2003; Malhi et al., 2008). The Amazon basin has a large vertebrate biodiversity (Silva et al., 2005). For example, the total number of freshwater fish species present in the Amazon basin represents ~15% of all freshwater fishes described worldwide (Jézéquel et al., 2020). Similarly, for three groups of terrestrial vertebrates (birds, mammals and amphibians), the Brazilian Amazon has a higher overall species richness compared with other Brazilian biomes (Jenkins et al., 2015). Vertebrates have great importance in the management of tropical forest ecosystems (Janzen, 1970). This includes seed dispersal, predation, regulation of water quality, and nutrient and carbon cycles in both terrestrial and aquatic ecosystems (Böhm et al., 2013; Fletcher et al., 2006; Raxworthy et al., 2008).

Amazon biodiversity is increasingly threatened by several factors, including habitat loss and fragmentation and climate change (Dudgeon et al., 2006; Laurance et al., 2011; Li et al., 2013; Malhi et al., 2008; Michalski & Peres, 2007; Schneider et al., 2021). One of the major threats to Amazonian biodiversity identified by the International Union for Conservation of Nature is the construction of hydroelectric power plants (IUCN, 2020). These constructions make a direct impact on the local environment and an indirect impact on a large scale, extending through the entire hydrology basin that is inserted (Carvalho et al., 2018). Expansion of hydropower developments in the Brazilian Amazon started in the 1980s (Fearnside, 2001; Junk et al., 1981), but only since 1986 does Brazilian legislation requires that developers need to produce a mandatory Environmental Impact Assessment (EIA), that evaluates the impact of the project and provides necessary minimization and mitigation actions. Although millions of dollars were invested, these EIAs are widely criticized as overly simplistic and generalist (Fearnside, 2014; Gerlak et al., 2020; Simões et al., 2014).

Systematic reviews summarize and evaluate studies, making evidence available for decision-makers (Gopalakrishnan & Ganeshkumar, 2013). A number of reviews document impacts of dams across the Amazon (Athayde, Mathews, et al., 2019; Ferreira et al., 2014; Lees et al., 2016). Recently several studies evaluated the impacts of hydroelectrics on water flow, sediments, and on aquatic Amazonian species, mostly fishes (Athayde, Mathews, et al., 2019; Castello et al., 2013; Latrubesse et al., 2017; Turgeon et al., 2021). But these and other reviews did not evaluate the quality of evidence presented in the primary studies. Indeed, to date there have been no systematic reviews on the impacts of hydroelectrics on Amazonian vertebrates.

In this review, we evaluated the scientific literature reporting hydroelectric impacts on vertebrates in Brazilian Amazonia. Specifically we addressed the following questions: (1) what are the temporal and spatial patterns of articles, (2) study designs adopted and (3) evidence types generated.

## Methods

### Study identification and selection

We focused on vertebrates as this group includes fish which is perhaps the most intensively studied wildlife group in terms of hydropower impacts globally (Algera et al., 2020; Arantes et al., 2019; Turgeon et al., 2021). Additionally, this group also includes “mega-fauna” (vertebrates > 30 kg) that have a disproportionately high risk of extinction due to human threats (He et al., 2018). As such vertebrates should present a best case scenario for the scientific evidence documenting hydropower impacts on threatened Amazonian wildlife. Searches were conducted for articles published from 1945 to 2020 using four different databases: ISI Web of Science, SCOPUS, PubMed and Scielo. The databases were searched using the following combination of terms: (Amazon*) and (hydroelectric or hydropower or dam) and (mammal or fish or bird or reptile or amphibian or vertebrate) and (impact* or effect*). The same terms were translated and searches repeated in Portuguese and Spanish. Searches were conducted twice, once on 28 March 2020 and again on 29 March 2021 to update publications from 2020.

Studies were selected following guidelines established by the Preferred Reporting Items for a Systematic Review and Meta-analysis [PRISMA (Moher et al., 2015; Shamseer et al., 2015), Figure 1]. First, we screened all titles, keywords and abstracts and excluded duplicates and any studies that were not related to hydroelectric developments and vertebrates within the legal Brazilian Amazon. The full-text of all articles that passed initial screening was then read to establish eligibility.

**Figure 1.**
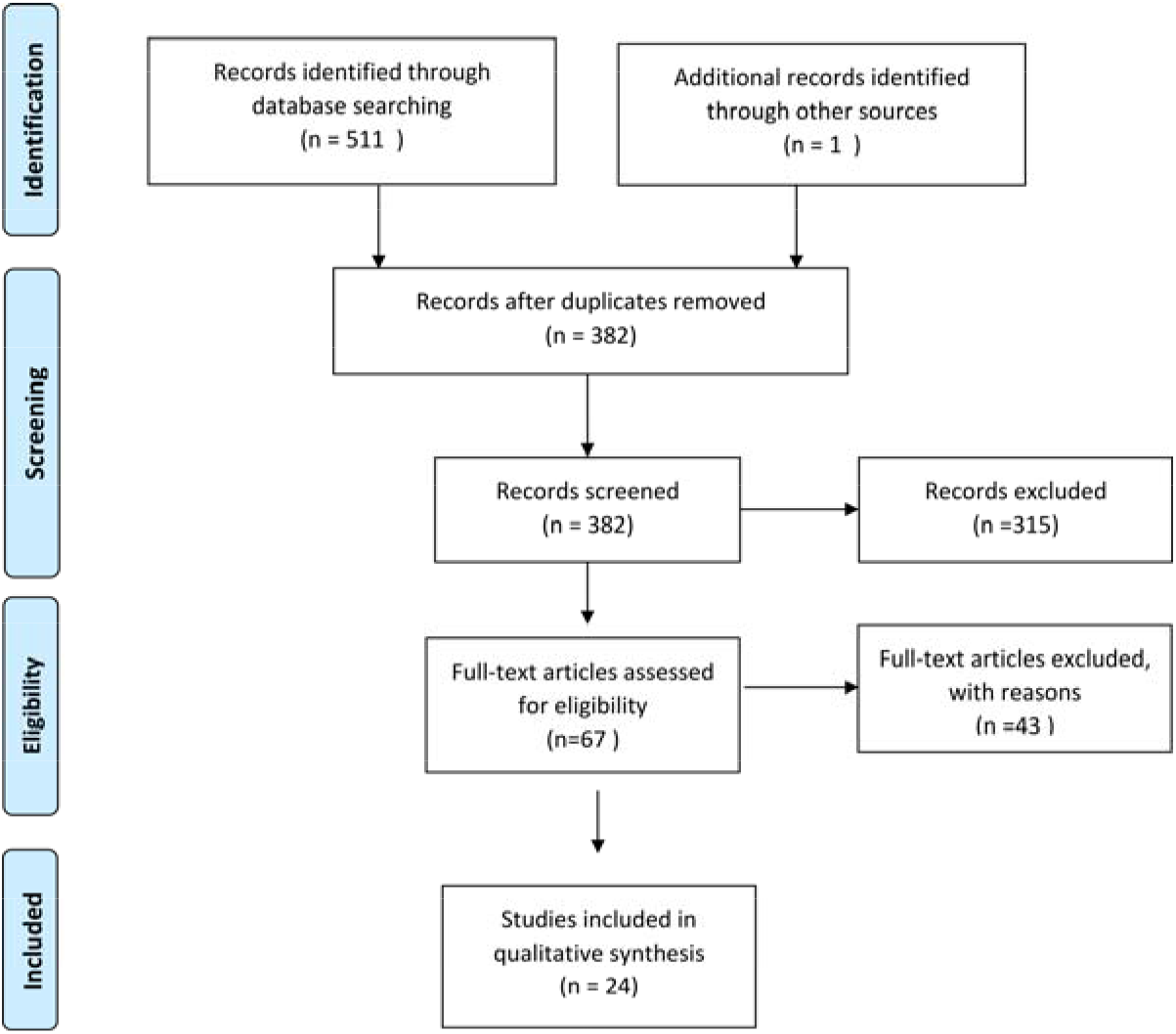
PRISMA flow chart. Showing process used to assess and select studies.

As our focus was on evaluating impacts, the studies needed to include results from comparisons with at least one of the following: control areas (including space-for-time) and/or the impacted area after the hydroelectric was operational. Selected articles needed to present basic data/primary studies (Salafsky et al., 2019) from operational hydroelectrics, as such laboratory experiments, simulations, reviews and meta-analysis were not included. Studies that used novel reservoir environments to test theories (e.g. species-area relationships on reservoir islands) were not included. In addition, studies with lists of species compared with other areas in only a qualitative narrative form or where comparisons were only discussed (not included as part of the sampling methodology or analysis) were also excluded at this stage.

### Study data extraction

Each study was evaluated by one reviewer, who compiled: publication year, vertebrate groups, period of data collection, study design, geographic coordinates for the studied dams [obtained by joining dam name with coordinates provided by SIGEL (2021)], evidence type and whether the study received funding/data from the developer/operator (Supplemental Material Appendix 1). Study design typology followed definitions in Christie et al. (2019) and evidence types were classified following Burivalova et al. (2019) (Table 1). Finally the PRISMA process and data extraction stages were independently reviewed by two researchers (DN and FM) and corrections made to ensure reproducibility and consistency.

**Table 1.**
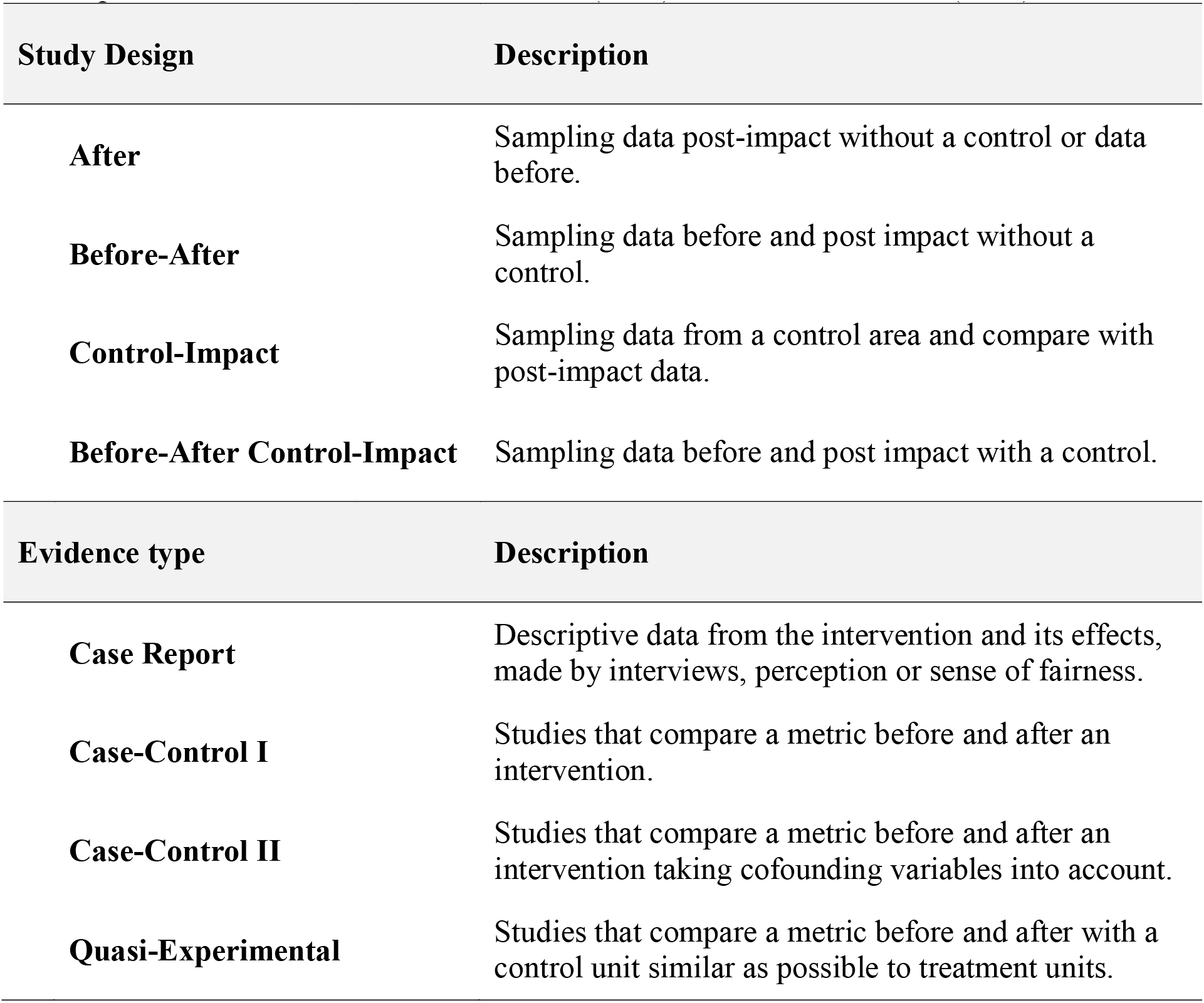
Study Designs and Evidence Types. Typology used to classify selected studies. Descriptions summarized from Christie et al. (2019) and Burivalova et al. (2019).

### Hydroelectric data

To contextualize the literature review we compiled data on the operational hydroelectric plants in the legal Brazilian Amazon. For each hydroelectric plant we obtained geographic coordinates, operational start date and power output from the Brazilian Electric Sector Geographic Information (SIGEL – “Sistema de Informações Georreferenciadas do Setor Elétrico”), provided and maintained by the Brazilian National Agency of Electricity (ANEEL – “Agência Nacional de Energia Elétrica”, downloaded from: https://sigel.aneel.gov.br/Down/, accessed on 30 March 2021). We retained only hydroelectric power plants (HPPs) with an installed power greater than 30 MW (Supplemental Material Appendix 2). We used ArcGIS 10.3 (ESRI, 2015) in order to produce the final distribution map of the hydroelectric plants and study locations.

### Data Analysis

All analyses were performed in R (R Development Core Team, 2020). Patterns in the geographic and temporal distribution of publications were evaluated using maps and descriptive analysis. As Brazilian states are an important administrative and legislative unit for the management of environmental resources, we compared the distribution of hydroelectrics and publications between the nine states of the 5 Mkm^2^ Legal Brazilian Amazon [Acre, Amapá, Amazonas, Mato Grosso, Maranhão, Pará, Rondônia, Roraima and Tocantins, (IBGE, 2020)]. The distribution of study designs and evidence types was compared between studies that i) received funding and/or data from the hydroelectric developer/operator and ii) independent research studies without any declared association with the hydroelectric developer/operator.

## Results

### Temporal and spatial distribution of studies

A total of 24 peer-reviewed studies were included in our review most of which (n = 16) were published between 2015 and 2020 (Figure 2). The first article found in our review was published in 1981 (Junk et al., 1981). This was four years after the hydroelectric plant under study (“Curuá-Una”) became operational in 1977 and six years after the first hydroelectric plant became operational in the legal Brazilian Amazon in 1975 (Figure 2). Although the number of operational hydroelectrics increased steadily in the subsequent decades, the number of published articles started to increase only recently (Figure 2). After the first published study there was a 12 year gap until the next publication and few studies (n = 4) were published by 2012, despite there being 15 operational hydroelectrics in 2010.

**Figure 2.**
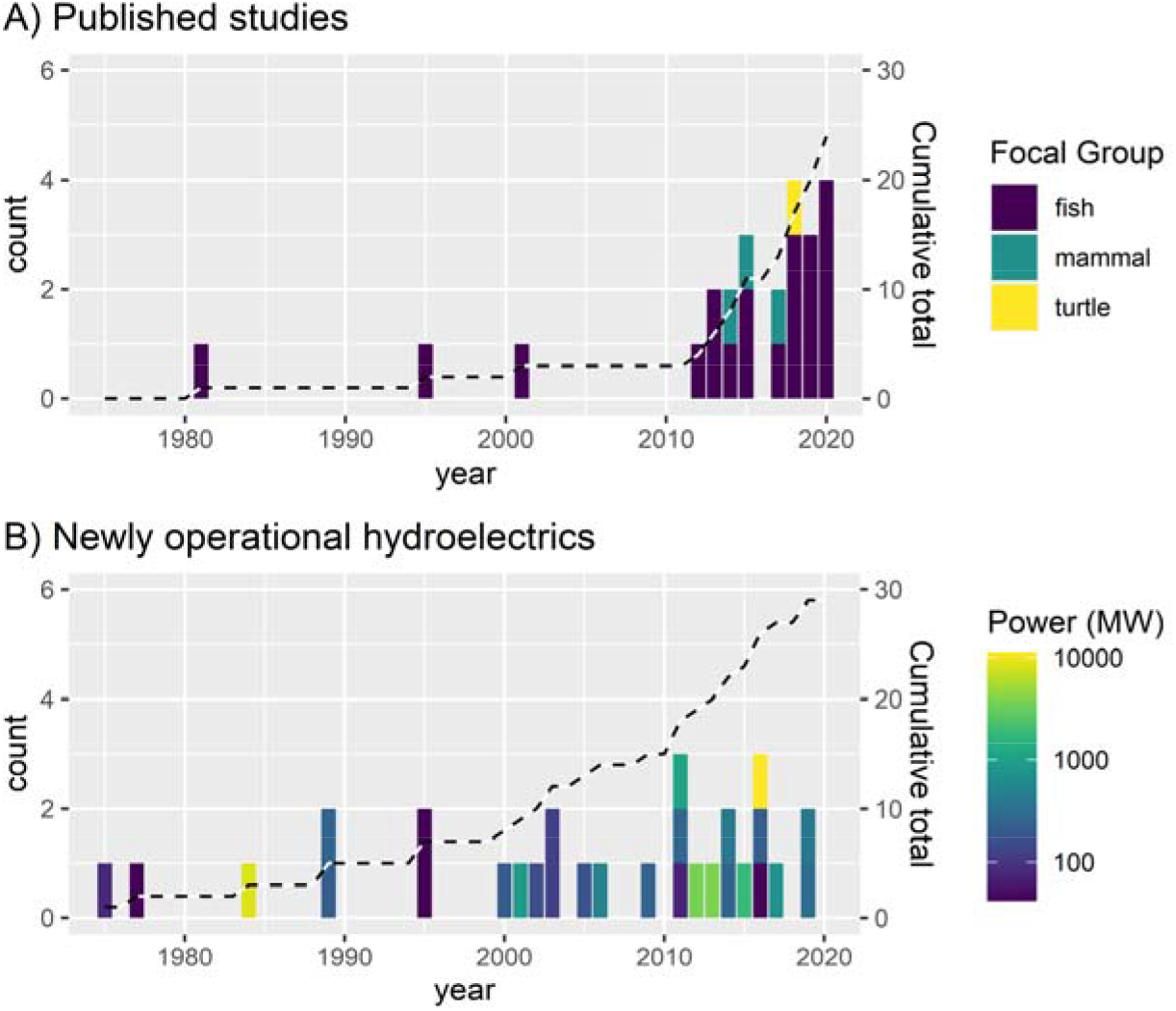
Temporal distribution of published studies and operational hydroelectrics. Annual frequency of A) published articles documenting impacts on vertebrates (n = 24) and B) newly operational hydroelectrics (n = 29) across the legal Brazilian Amazon. Dashed lines show cumulative totals.

Based on our inclusion criteria we were able to identify studies assessing impacts on only three groups of vertebrates (Figure 2): fish (n = 20), mammals (n = 3) and turtles (n = 1). The major research interest was related to fish (83.3% of studies) with the four articles published during the first three decades (1981 – 2013) focusing exclusively on this group (Figure 2). The three mammal studies (Calaça & de Melo, 2017; Calaça et al., 2015; Palmeirim et al., 2014) were published between 2014 and 2017 and all focused on the semi-aquatic Giant Otter (*Pteronura brasiliensis*). The study assessing impacts on turtles (Norris et al., 2018) focused on the Yellow-spotted River Turtle (*Podocnemis unifilis*).

The studies assessed impacts caused by 12 of the 29 operational hydroelectric plants. The distribution of studies tended to follow the power output of the dams in each state (Figure 3) and we found a positive but insignificant correlation between power output and number of studies per hydroelectric power plant (Spearman Correlation rho = 0.41, p = 0.181). Nearly half of studies (n= 11) investigated impacts of three power plants, namely Jirau and Santo Antônio (n = 7, with 6 studies including both) in the state of Rondônia and Peixe Angical (n = 4) in Tocantins. With the two most intensely studied hydroelectrics (Jirau and Santo Antonio, power output 3750 and 3568 MW respectively) accounting for 7 of the 13 studies published since 2017. The remaining 9 hydroelectric plants had one or two studies each. We also found a weak positive correlation between the number of hydroelectrics and number of published studies per state (Spearman Correlation rho = 0.21, p = 0.686). Mato Grosso was the state with most hydroelectric power plants (n = 13), but was severely under-represented with only two published studies (Figure 3), both of which focused around the recently operational Teles Pires dam [1,819 MW, operational in November 2015, (Calaça & de Melo, 2017; Calaça et al., 2015)].

**Figure 3.**
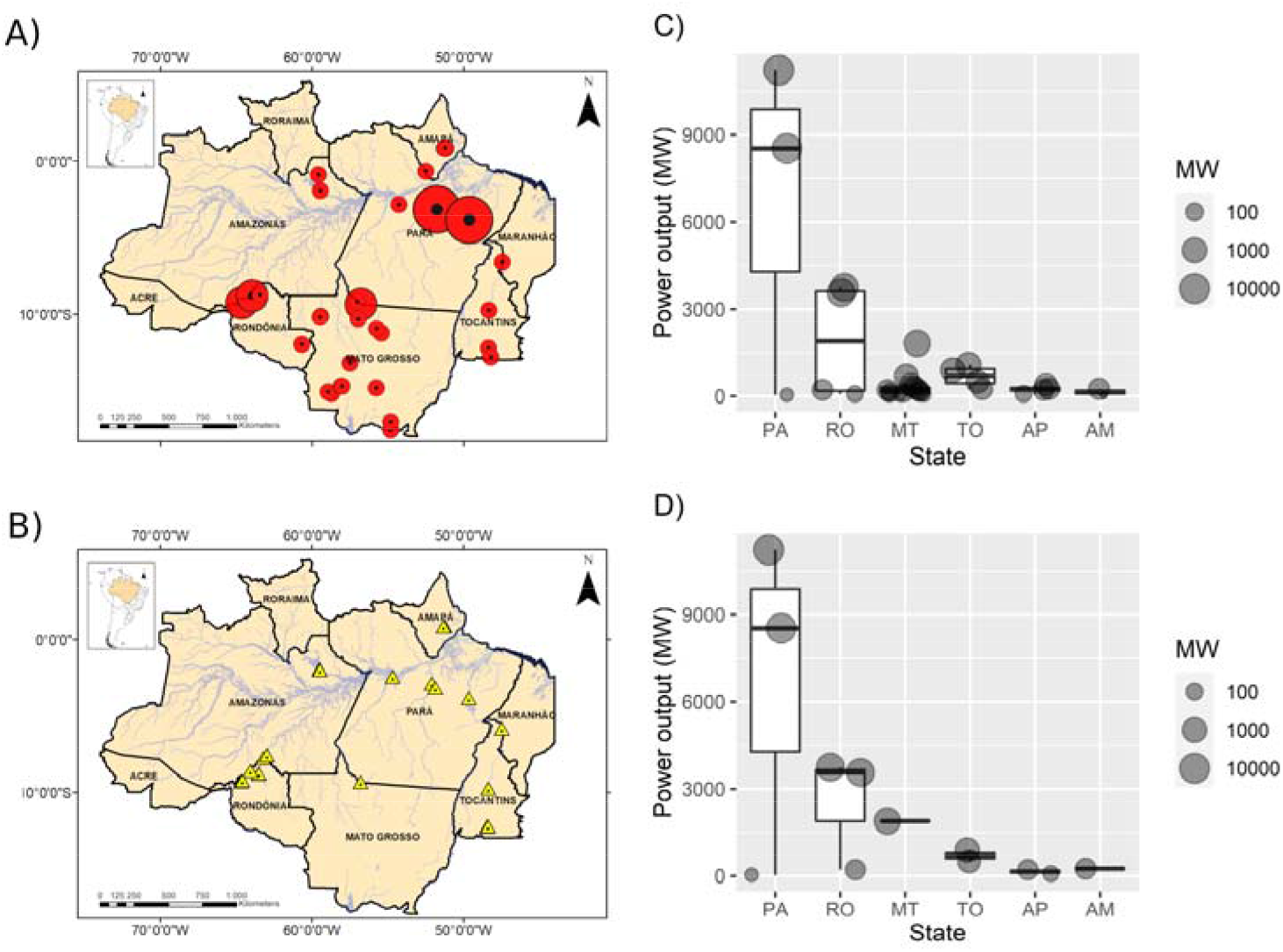
Spatial distribution of published studies and operational hydroelectrics. Geographic location of A) operational hydroelectrics (circles, n = 29) and B) studies documenting impacts on vertebrates (triangles, n = 24) across the legal Brazilian Amazon. The size of the circles showing hydroelectric locations is proportional to the power output of each hydroelectric, and light grey lines represent major rivers. Plots show distribution of power output (MW) by C) State of all 29 operational hydroelectric and D) The 12 hydroelectrics included in 24 studies. The sequence of States is ordered by total power output of operational hydroelectrics in each state (high to low from left to right).

### Study Design and Evidence Type

Most studies (87.5%) adopted either “After” (n = 6) or “Before-After” (n = 15) study designs (Figure 4). Only three studies used a Before-After Control-Impact design, two with fish (Araújo et al., 2013; Lima et al., 2018) and one with turtles (Norris et al., 2018).

**Figure 4.**
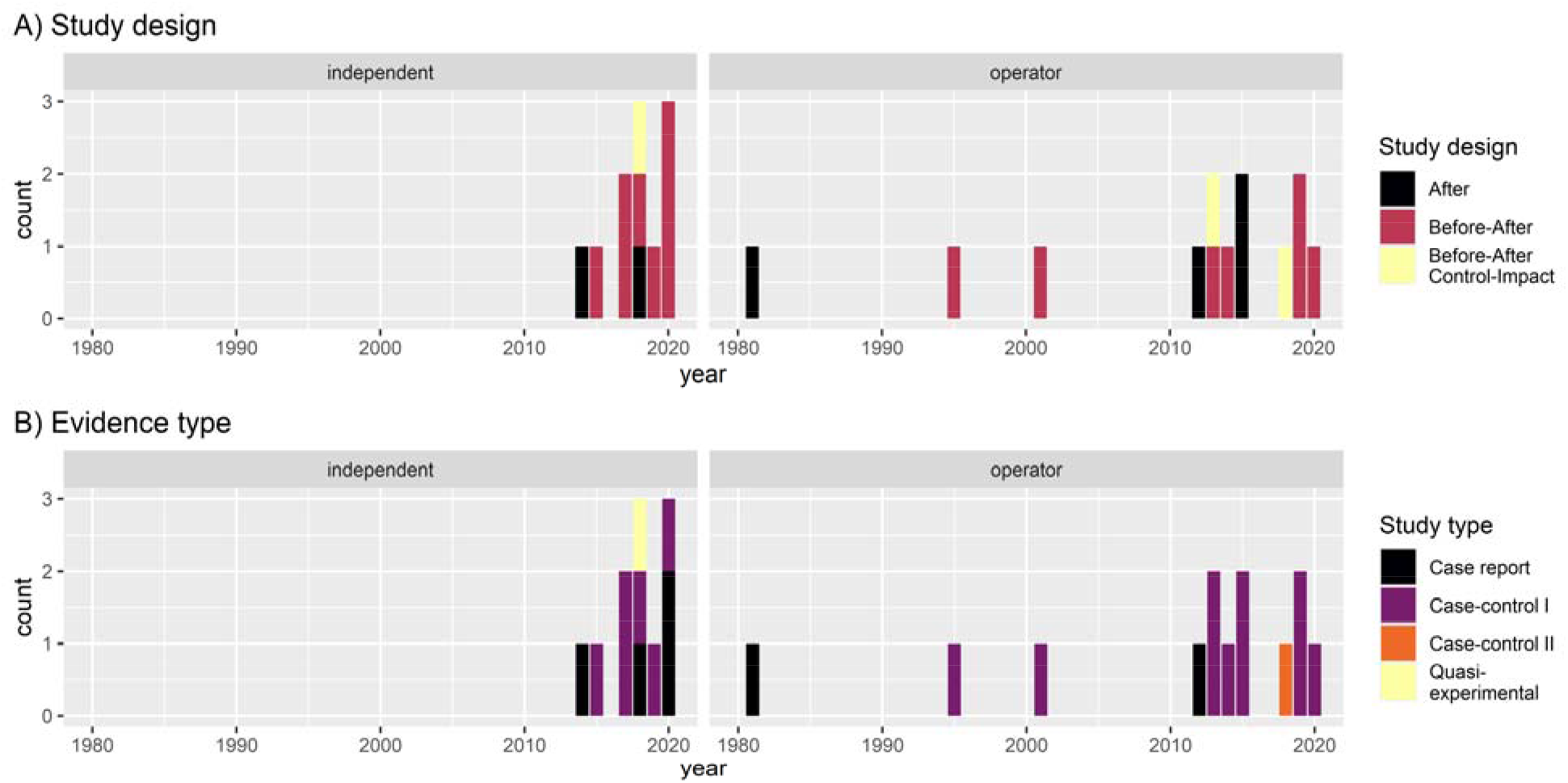
Temporal distribution of study designs and evidence types. The A) study design used and B) type of evidence produced by 24 published articles documenting impacts hydroelectric developments on vertebrates across the legal Brazilian Amazon. Classification follows previously published definitions of study designs (Christie et al., 2019) and evidence types (Burivalova et al., 2019). Studies are grouped into those conducted without financial support from the developer/operator (“independent”) and those that received financial support or data from the developer/operator (“operator”).

Most publications (91.7%, n = 22) did not support causal inference, with evidence coming from either Case-report (n = 6) or Case-Control I (n = 16) studies (Figure 4). Only one Quasi-Experimental study was found, which included data collected pre and post reservoir formation with both impacted and control areas and analysis to explicitly test the Before-After Control-Impact interaction (Norris et al., 2018). The proportion of independent (n = 11) and operator funded (n = 13) studies was similar (Chi-squared = 0.17, df = 1, p = 0.683) and there was no significant difference in the frequencies of study designs or evidence types between independently or operator funded studies (Figure 4, Fisher’s Exact Test p = 0.725 and 0.288 for study designs and evidence types respectively).

## Discussion

Our systematic review showed that (1) studies focused on understanding the impacts of hydroelectrics on Amazonian vertebrates are increasing, but weak sampling designs resulted in a lack of robust evidence, (2) the majority of studies focused on fish, and (3) there was a tendency for studies to be concentrated on high potency “mega” hydropower plants. We first turn to discuss the lack of evidence due to weak sampling designs and then explore the focus on selected vertebrate groups, discrepancy on studies focused on large dams and lack of integrated studies.

The lack of robust evidence was surprising considering hydropower development impacts are so strong and well known at a global scale (Grill et al., 2019; Liermann et al., 2012; Maavara et al., 2020). We found that studies across Brazilian Amazonia were biased by a focus on mega-dams. A major part of the increasing number of studies since 2012 can be attributed to studies of only two dams (Jirau and Santo Antonio). Although the sustainability of both projects was questioned (Fearnside, 2014, 2015), both received certification by Hydropower Sustainability Assessment Protocol (https://www.hydrosustainability.org/published-assessments/santo-antonio and https://www.hydrosustainability.org/published-assessments/jirau, accessed 23 June 2021). Our results show that scientific evidence documenting the impacts of both was generally weak (i.e. below expected best practice). A finding that supports recent analysis showing a link between superficial impact assessments and a lack of social and environmental sustainability of Amazonian hydropower developments (Fearnside, 2018; Gerlak et al., 2020).

We found that studies generally adopted weak sampling designs (e.g. lacking controls) and lacked evidence necessary to generate reliable inference (Christie et al., 2021; Christie et al., 2019; Salafsky et al., 2019). Although randomized-control studies are widely recognized as the most robust, logistically simpler designs such as before-after control-impact can be equally effective in generating robust evidence for impact assessments of abrupt changes induced by large scale development projects including dam construction. Additionally, dams are so widespread across Amazonia (Anderson et al., 2018; Athayde, Duarte, et al., 2019; Grill et al., 2019) that there are few remaining free flowing river sections that could be included within a randomized-control design.

Most of the studies found in our review focused on fishes and are therefore likely to represent best-case scenario in terms of scientific knowledge and evidence base. In fact, this finding follows global patterns where fishes were one of the most frequently studied groups used to evaluate effects of hydroelectric dams in both temperate (Algera et al., 2020) and tropical regions (Arantes et al., 2019). But, impacts of run-of-river dams are poorly studied even for fish the most intensively studied group (Turgeon et al., 2021). Moreover, there is a lack of studies on multiple vertebrate groups, which is essential to understand hydroelectric effects on complex hydrological systems such as the Amazon (Park & Latrubesse, 2017).

As impacts are so poorly understood it is also unsurprising that there is limited evidence documenting the effectiveness of mitigation actions for vertebrates impacted by hydropower developments across Amazonia. For example, from a total of 48 actions identified in the Conservation Evidence database (https://www.conservationevidence.com/data/index?pp=50&terms=dam&country%5B%5D=&result_type=interventions&sort=relevance.desc#searchcontainer, accessed 14 July 2021) there were no studies from the Amazon basin. Although it is possible to suggest some general actions based on documented global experiences, no studies have evaluated effects of installing bypasses channels for aquatic mammals (Berthinussen et al., 2021) and only three short-term studies (10 to 18 months) evaluated translocations, two in French Guiana, both for primates (Richard-Hansen et al., 2000; Vié et al., 2001) and one in central Brazil for lesser anteater (Rodrigues et al., 2009). Indeed, to date no studies have implemented or evaluated mitigation actions that are likely to generate multiple conservation benefits such as habitat restoration.

Our review showed a lack of studies assessing multiple hydroelectrics and/or multiple vertebrate groups along the same river. In Brazil, several hydroelectric plants belonging to different operators are commonly arranged in the same river, creating “cascades” (Athayde, Duarte, et al., 2019; Mendes et al., 2017). Although many studies focus on mega-dams, the combined effect of multiple hydroelectrics, which can cause cumulative impacts (Athayde, Duarte, et al., 2019) remains poorly documented. For example, Coaracy Nunes was the first dam installed in the legal Brazilian Amazon in 1975, since then two additional dams have become operational along the same river, providing a total of three dams with a combined output of 549 MW (78, 252 and 219 MW) within a 18 km stretch of river. The impact of these multiple dams is thought to have drastically altered both upstream and downstream flow rates and following the installation of the second dam (Ferreira Gomes) in 2014 the rivers downstream course became divided, draining predominantly to the Amazon river not the Atlantic Ocean (Silva dos Santos, 2017). Whilst individual studies focus on fish (Sá-Oliveira et al., 2015; Sá-Oliveira et al., 2016) and turtles (Norris et al., 2018; Norris et al., 2020) along the impacted river, these studies focused on different dams and adopted different sampling designs, which limits the ability to integrate results for important basin wide analysis necessary to inform mitigation actions.

We failed to find studies including important cofounding impacts such as deforestation (Stickler et al., 2013). Although deforestation and tree mortality have been widely documented as important impacts of Amazonian dams (Athayde, Mathews, et al., 2019; Resende et al., 2019; Stickler et al., 2013) no studies included these important cofounding variables in the assessments of vertebrates. For example, the lack of studies in Mato Grosso was particularly surprising considering previous studies on effects of forest fragmentation on vertebrates in this state (Michalski & Peres, 2007; Norris & Michalski, 2009).

We found few studies considering the overall number and investment in hydropower projects across the Legal Brazilian Amazonia. Even fewer studies were found when considering only those with a robust design and able to establish causal inference. It could be suggested that weak evidence is a reflection of a lack of investment in science and technology, together with a reduction in investment in the Brazilian Ministry of the Environment over the past twenty years (de Area Leão Pereira et al., 2019). Although there is undoubtedly support for such considerations, the lack of robust survey designs can also perhaps be attributed more simply to a failure of researchers to adopt robust designs (Christie et al., 2021; Christie et al., 2019).

However, we need to highlight that our review has some limitations, as we did not include “grey literature” in our searchers. Thus, it is important to recognize the potential for gaps or missing studies that were not published in peer-reviewed journals. On the other hand, as we would expect published studies to have more robust designs and analysis compared with grey literature or reports, our review, performed in searches across four different databases and in three languages is likely to be a best-case representation of the scientific evidence base documenting hydroelectric impacts on vertebrates in the Brazilian Amazonia.

### Implications for conservation

There is an urgent need to take advantage of freely available data to organize and plan effective surveys and sampling strategies to evaluate sustainability of current and future hydroelectric across the Brazilian Amazon. Below we provide recommendations to help develop a more robust evidence base.

1. **Geographical distribution of studies.** **Research gaps**: Studies were focused within specific regions **Future directions**: Increase the number of studies all around Brazilian Amazon with a focus in Mato Grosso state, which has more than 50% of operational and planned hydroelectrics.
2. **Study groups.** **Research gaps**: The majority of studies focus on understanding the impacts on fish. **Future directions**: Increase studies focusing on other threatened vertebrate groups including amphibians, birds, mammals, and reptiles.
3. **Hydroelectric power plants.** **Research gaps**: Most of our reviewed studies were concentrated in three large hydroelectric power plants. **Future directions**: Increase number of studies to represent the distribution of operational and planned power output. This should include closer integration with university research teams to develop robust evidence as part of the necessary Environmental Impact Assessments.
4. **Study design and evidence.** **Research gaps**: There is currently a lack of robust evidence to evaluate impacts of hydroelectric power plants on Amazonian wildlife. **Future directions**: Studies need to include more robust designs (e.g. Before-After Control-Impact) to establish causal inference.

## Acknowledgements

The Federal University of Amapá (UNIFAP) provided logistical support.

## Referrences

Algera, D. A., Rytwinski, T., Taylor, J. J., Bennett, J. R., Smokorowski, K. E., Harrison, P. M., … Cooke, S. J. (2020). What are the relative risks of mortality and injury for fish during downstream passage at hydroelectric dams in temperate regions? A systematic review. Environmental Evidence, 9(1), 3. doi:10.1186/s13750-020-0184-0

Anderson, E. P., Jenkins, C. N., Heilpern, S., Maldonado-Ocampo, J. A., Carvajal-Vallejos, F. M., Encalada, A. C., … Tedesco, P. A. (2018). Fragmentation of Andes-to-Amazon connectivity by hydropower dams. Sci Adv, 4(1), eaao1642. doi:10.1126/sciadv.aao1642

Arantes, C. C., Fitzgerald, D. B., Hoeinghaus, D. J., & Winemiller, K. O. (2019). Impacts of hydroelectric dams on fishes and fisheries in tropical rivers through the lens of functional traits. Current Opinion in Environmental Sustainability, 37, 28–40. doi:https://doi.org/10.1016/j.cosust.2019.04.009

Araújo, E. S., Marques, E. E., Freitas, I. S., Neuberger, A. L., Fernandes, R., & Pelicice, F. M. (2013). Changes in distance decay relationships after river regulation: similarity among fish assemblages in a large Amazonian river. Ecology of Freshwater Fish, 22(4), 543–552. doi:https://doi.org/10.1111/eff.12054

Athayde, S., Duarte, C. G., Gallardo, A. L. C. F., Moretto, E. M., Sangoi, L. A., Dibo, A. P. A., … Sánchez, L. E. (2019). Improving policies and instruments to address cumulative impacts of small hydropower in the Amazon. Energy Policy, 132, 265–271. doi:https://doi.org/10.1016/j.enpol.2019.05.003

Athayde, S., Mathews, M., Bohlman, S., Brasil, W., Doria, C., Dutka-Gianelli, J., … Kaplan, D. (2019). Mapping research on hydropower and sustainability in the Brazilian Amazon: advances, gaps in knowledge and future directions. Current Opinion in Environmental Sustainability, 37, 50–69. doi:https://doi.org/10.1016/j.cosust.2019.06.004

Aurelio-Silva, M., Anciaes, M., Henriques, L. M. P., Benchimol, M., & Peres, C. A. (2016). Patterns of local extinction in an Amazonian archipelagic avifauna following 25 years of insularization. Biological Conservation, 199, 101–109. doi:https://doi.org/10.1016/j.biocon.2016.03.016

Benchimol, M., & Peres, C. A. (2015). Widespread Forest Vertebrate Extinctions Induced by a Mega Hydroelectric Dam in Lowland Amazonia. Plos One, 10(7). doi:https://doi.org/10.1371/journal.pone.0129818

Berthinussen, A., Smith, R. K., & Sutherland, W. J. (2021). Marine and Freshwater Mammal Conservation: Global Evidence for the Effects of Interventions. Conservation Evidence Series Synopses. Retrieved from https://www.conservationevidence.com/synopsis/pdf/30

Böhm, M., Collen, B., Baillie, J., Bowles, P., Chanson, J., Cox, N., … Zug, G. (2013). The Conservation Status of the World’s Reptiles. Biological Conservation, 157, 372–385. doi:https://doi.org/10.1016/j.biocon.2012.07.015

Bueno, A. S., & Peres, C. A. (2019). Patch-scale biodiversity retention in fragmented landscapes: Reconciling the habitat amount hypothesis with the island biogeography theory. Journal of Biogeography, 46(3), 621–632. doi:https://doi.org/10.1111/jbi.13499

Burivalova, Z., Miteva, D., Salafsky, N., Butler, R. A., & Wilcove, D. S. (2019). Evidence Types and Trends in Tropical Forest Conservation Literature. Trends in Ecology & Evolution, 34(7), 669–679. doi:https://doi.org/10.1016/j.tree.2019.03.002

Calaça, A. M., & de Melo, F. R. (2017). Reestablishment of giant otters in habitats altered by the filling of the Teles Pires hydroelectric dam in the Amazonia. IUCN Otter Specialist Group Bulletin, 34(2), 73–78.

Calaça, A. M., Faedo, O. J., & de Melo, F. R. (2015). Hydroelectric Dams: The First Responses from Giant Otters to a Changing Environment. IUCN Otter Spec. Group Bull, 32(1), 48–58.

Carvalho, D. N., Boniolo, M. R., Santos, R. G., Batista, L. V., Malavazzi, A. A., Reis, F. A. G. V., & Giordano, L. d. C. (2018). Criteria applied in the definition of influence areas, impacts and programmes in environmental impact studies of Brazilian hydroelectric power plants. Geociencias - UNESP, 37, 15.

Castello, L., McGrath, D. G., Hess, L. L., Coe, M. T., Lefebvre, P. A., Petry, P., … Arantes, C. C. (2013). The vulnerability of Amazon freshwater ecosystems. Conservation Letters, 6(4), 217–229. doi:https://doi.org/10.1111/conl.12008

Christie, A. P., Amano, T., Martin, P. A., Petrovan, S. O., Shackelford, G. E., Simmons, B. I., … Sutherland, W. J. (2021). The challenge of biased evidence in conservation. Conservation Biology, 35(1), 249–262. doi:https://doi.org/10.1111/cobi.13577

Christie, A. P., Amano, T., Martin, P. A., Shackelford, G. E., Simmons, B. I., & Sutherland, W. J. (2019). Simple study designs in ecology produce inaccurate estimates of biodiversity responses. Journal of Applied Ecology, 56(12), 2742–2754. doi:https://doi.org/10.1111/1365-2664.13499

Cosson, J. F., Ringuet, S., Claessens, O., de Massary, J. C., Dalecky, A., Villiers, J. F., … Pons, J. M. (1999). Ecological changes in recent land-bridge islands in French Guiana, with emphasis on vertebrate communities. Biological Conservation, 91(2-3), 213–222. doi:https://doi.org/10.1016/s0006-3207(99)00091-9

de Area Leão Pereira, E. J., Silveira Ferreira, P. J., de Santana Ribeiro, L. C., Sabadini Carvalho, T., & de Barros Pereira, H. B. (2019). Policy in Brazil (2016–2019) threaten conservation of the Amazon rainforest. Environmental Science & Policy, 100, 8–12. doi:https://doi.org/10.1016/j.envsci.2019.06.001

Dirzo, R., & Raven, P. H. (2003). Global State of Biodiversity and Loss. Annual Review of Environment and Resources, 28(1), 137–167. doi:https://doi.org/10.1146/annurev.energy.28.050302.105532

Dudgeon, D., Arthington, A. H., Gessner, M. O., Kawabata, Z. I., Knowler, D. J., Leveque, C., … Sullivan, C. A. (2006). Freshwater biodiversity: importance, threats, status and conservation challenges. Biological Reviews, 81(2), 163–182. doi:https://doi.org/10.1017/s1464793105006950

Egré, D., & Milewski, J. C. (2002). The diversity of hydropower projects. Energy Policy, 30(14), 1225–1230. doi:https://doi.org/10.1016/S0301-4215(02)00083-6

ESRI. (2015). ArcGIS Desktop: Release 10.3. Redlands, CA: Environmental Systems Research Institute.

Fearnside, P. M. (1989). Brazil’s Balbina Dam: Environment versus the legacy of the Pharaohs in Amazonia. Environ Manage, 13(4), 401–423. doi:https://doi.org/10.1007/BF01867675

Fearnside, P. M. (2001). Environmental impacts of Brazil’s Tucurui Dam: unlearned lessons for hydroelectric development in Amazonia. Environ Manage, 27(3), 377–396. doi:https://doi.org/10.1007/s002670010156

Fearnside, P. M. (2006). Dams in the Amazon: Belo Monte and Brazil’s Hydroelectric Development of the Xingu River Basin. Environ Manage, 38(1), 16. doi:https://doi.org/10.1007/s00267-005-0113-6

Fearnside, P. M. (2014). Impacts of Brazil’s Madeira River Dams: Unlearned lessons for hydroelectric development in Amazonia. Environmental Science & Policy, 38, 164–172. doi:https://doi.org/10.1016/j.envsci.2013.11.004

Fearnside, P. M. (2015). Tropical hydropower in the clean development mechanism: Brazil’s Santo Antônio Dam as an example of the need for change. Climatic Change, 131(4), 575–589. doi:https://doi.org/10.1007/s10584-015-1393-3

Fearnside, P. M. (2018). Challenges for sustainable development in Brazilian Amazonia. Sustainable Development, 26(2), 141–149. doi:https://doi.org/10.1002/sd.1725

Ferreira, J., Aragão, L. E. O. C., Barlow, J., Barreto, P., Berenguer, E., Bustamante, M., … Zuanon, J. (2014). Brazil’s environmental leadership at risk. Science, 346(6210), 706. doi:https://doi.org/10.1126/science.1260194

Finer, M., & Jenkins, C. N. (2012). Proliferation of hydroelectric dams in the Andean Amazon and implications for Andes-Amazon connectivity. Plos One, 7(4), e35126–e35126. doi:https://doi.org/10.1371/journal.pone.0035126

Fletcher, D., Hopkins, W., Saldaña, T., Baionno, J., Arribas, C., Standora, M., & Fernandez-Delgado, C. (2006). Geckos as indicators of mining pollution. Environmental toxicology and chemistry / SETAC, 25, 2432–2445. doi:https://doi.org/10.1897/05-556R.1

Gerlak, A. K., Saguier, M., Mills-Novoa, M., Fearnside, P. M., & Albrecht, T. R. (2020). Dams, Chinese investments, and EIAs: A race to the bottom in South America? Ambio, 49(1), 156–164. doi:https://doi.org/10.1007/s13280-018-01145-y

Gopalakrishnan, S., & Ganeshkumar, P. (2013). Systematic reviews and meta-analysis: Understanding the best evidence in primary healthcare. 2(1), 9–14. doi:https://doi.org/10.4103/2249-4863.109934

Grill, G., Lehner, B., Thieme, M., Geenen, B., Tickner, D., Antonelli, F., … Zarfl, C. (2019). Mapping the world’s free-flowing rivers. Nature, 569(7755), 215–221. doi:https://doi.org/10.1038/s41586-019-1111-9

Hall, A., & Branford, S. (2012). Development, Dams and Dilma: the Saga of Belo Monte. Critical Sociology, 38(6), 851–862. doi:https://doi.org/10.1177/0896920512440712

He, F., Bremerich, V., Zarfl, C., Geldmann, J., Langhans, S. D., David, J. N. W., … Jähnig, S. C. (2018). Freshwater megafauna diversity: patterns, status and threats. Diversity Distributions, 24(10), 1395–1404. doi:https://doi.org/10.1111/ddi.12780

IBGE. (2020). Legal Amazon Boundaries for 2019 [Press release]. Retrieved from https://censos.ibge.gov.br/en/2185-news-agency/releases-en/28109-ibge-updates-map-of-the-legal-amazon.html

IUCN. (2020). The IUCN Red List of Threatened Species.

Janzen, D. H. (1970). Herbivores and the Number of Tree Species in Tropical Forests. 104(940), 501–528. doi:https://doi.org/10.1086/282687

Jenkins, C. N., Alves, M. A. S., Uezu, A., & Vale, M. M. (2015). Patterns of Vertebrate Diversity and Protection in Brazil. Plos One, 10(12), e0145064. doi:https://doi.org/10.1371/journal.pone.0145064

Jézéquel, C., Tedesco, P. A., Bigorne, R., Maldonado-Ocampo, J. A., Ortega, H., Hidalgo, M., … Oberdorff, T. (2020). A database of freshwater fish species of the Amazon Basin. Scientific Data, 7(1), 96. doi:https://doi.org/10.1038/s41597-020-0436-4

Junk, W. J., Robertson, B. A., Darwich, A. J., & Vieira, I. (1981). Investigações limnológicas e ictiológicas em Curuá-Una, a primeira represa hidrelétrica na Amazônia Central. Acta Amazonica, 11(4), 689–717. doi:https://doi.org/10.1590/1809-43921981114689

Latrubesse, E. M., Arima, E. Y., Dunne, T., Park, E., Baker, V. R., d’Horta, F. M., … Stevaux, J. C. (2017). Damming the rivers of the Amazon basin. Nature, 546(7658), 363–369. doi:https://doi.org/10.1038/nature22333

Laurance, W. F., Camargo, J. L. C., Luizao, R. C. C., Laurance, S. G., Pimm, S. L., Bruna, E. M., … Lovejoy, T. E. (2011). The fate of Amazonian forest fragments: A 32-year investigation. Biological Conservation, 144(1), 56–67. doi:https://doi.org/10.1016/j.biocon.2010.09.021

Lees, A. C., Peres, C. A., Fearnside, P. M., Schneider, M., & Zuanon, J. A. S. (2016). Hydropower and the future of Amazonian biodiversity. Biodiversity and Conservation, 25(3), 451–466. doi:https://doi.org/10.1007/s10531-016-1072-3

Li, J. S., Lin, X., Chen, A. P., Peterson, T., Ma, K. P., Bertzky, M., … Poulter, B. (2013). Global Priority Conservation Areas in the Face of 21st Century Climate Change. Plos One, 8(1), 9. doi:https://doi.org/10.1371/journal.pone.0054839

Liermann, C. R., Nilsson, C., Robertson, J., & Ng, R. Y. (2012). Implications of Dam Obstruction for Global Freshwater Fish Diversity. BioScience, 62(6), 539–548. doi:10.1525/bio.2012.62.6.5

Lima, A. C., Sayanda, D., Agostinho, C. S., Machado, A. L., Soares, A. M. V. M., & Monaghan, K. A. (2018). Using a trait-based approach to measure the impact of dam closure in fish communities of a Neotropical River. Ecology of Freshwater Fish, 27(1), 408–420. doi:https://doi.org/10.1111/eff.12356

Maavara, T., Chen, Q., Van Meter, K., Brown, L. E., Zhang, J., Ni, J., & Zarfl, C. (2020). River dam impacts on biogeochemical cycling. Nature Reviews Earth & Environment, 1(2), 103–116. doi:https://doi.org/10.1038/s43017-019-0019-0

Malhi, Y., Roberts, J. T., Betts, R. A., Killeen, T. J., Li, W. H., & Nobre, C. A. (2008). Climate change, deforestation, and the fate of the Amazon. Science, 319(5860), 169–172. doi:https://doi.org/10.1126/science.1146961

Mendes, C. A. B., Beluco, A., & Canales, F. A. (2017). Some important uncertainties related to climate change in projections for the Brazilian hydropower expansion in the Amazon. Energy, 141, 123–138. doi:https://doi.org/10.1016/j.energy.2017.09.071

Mendes, Y. A., Oliveira, R. S., Montag, L. F. A., Andrade, M. C., Giarrizzo, T., Rocha, R. M., & Auxiliadora P. Ferreira, M. (2021). Sedentary fish as indicators of changes in the river flow rate after impoundment. Ecological Indicators, 125, 107466. doi:https://doi.org/10.1016/j.ecolind.2021.107466

Michalski, F., & Peres, C. A. (2007). Disturbance-Mediated Mammal Persistence and Abundance-Area Relationships in Amazonian Forest Fragments. Conservation Biology, 21(6), 1626–1640. doi:https://doi.org/10.1111/j.1523-1739.2007.00797.x

Moher, D., Shamseer, L., Clarke, M., Ghersi, D., Liberati, A., Petticrew, M., … Group, P.-P. (2015). Preferred reporting items for systematic review and meta-analysis protocols (PRISMA-P) 2015 statement. Systematic Reviews, 4(1), 1. doi:https://doi.org/10.1186/2046-4053-4-1

Norris, D., & Michalski, F. (2009). Are otters an effective flagship for the conservation of riparian corridors in an Amazon deforestation frontier. IUCN Otter Spec. Group Bull, 26(2), 72–76.

Norris, D., Michalski, F., & Gibbs, J. P. (2018). Beyond harm’s reach? Submersion of river turtle nesting areas and implications for restoration actions after Amazon hydropower development. PeerJ, 6, e4228. doi:https://doi.org/10.7717/peerj.4228

Norris, D., Michalski, F., & Gibbs, J. P. (2020). Community based actions save Yellow-spotted river turtle (Podocnemis unifilis) eggs and hatchlings flooded by rapid river level rises. PeerJ, 8, e9921. doi:10.7717/peerj.9921

Palmeirim, A. F., Peres, C. A., & Rosas, F. C. W. (2014). Giant otter population responses to habitat expansion and degradation induced by a mega hydroelectric dam. Biological Conservation, 174, 30–38. doi:https://doi.org/10.1016/j.biocon.2014.03.015

Palmeirim, A. F., Vieira, M. V., & Peres, C. A. (2017). Non-random lizard extinctions in land-bridge Amazonian forest islands after 28 years of isolation. Biological Conservation, 214, 55–65. doi:https://doi.org/10.1016/j.biocon.2017.08.002

Park, E., & Latrubesse, E. M. (2017). The hydro-geomorphologic complexity of the lower Amazon River floodplain and hydrological connectivity assessed by remote sensing and field control. Remote Sensing of Environment, 198, 321–332. doi:https://doi.org/10.1016/j.rse.2017.06.021

R Development Core Team. (2020). R: A language and enviroment for statistical computing. Vienna, Austria: R Fundation for Statistical Computing.

Raxworthy, C., Pearson, R., Zimkus, B., Reddy, S., Deo, A., Nussbaum, R., & Ingram, C. (2008). Continental speciation in the tropics: Contrasting biogeographic patterns of divergence in the Uroplatus leaf-tailed gecko radiation of Madagascar. Journal of Zoology, 275, 423–440. doi:https://doi.org/10.1111/j.1469-7998.2008.00460.x

Resende, A. F. d., Schöngart, J., Streher, A. S., Ferreira-Ferreira, J., Piedade, M. T. F., & Silva, T. S. F. (2019). Massive tree mortality from flood pulse disturbances in Amazonian floodplain forests: The collateral effects of hydropower production. Science of The Total Environment, 659, 587–598. doi:https://doi.org/10.1016/j.scitotenv.2018.12.208

Richard-Hansen, C., Vié, J. C., & de Thoisy, B. t. (2000). Translocation of red howler monkeys (Alouatta seniculus) in French Guiana. Biological Conservation, 93(2), 247–253. doi:https://doi.org/10.1016/S0006-3207(99)00136-6

Rodrigues, F. H. G., Marinho-Filho, J., & dos Santos, H. G. (2009). Home ranges of translocated lesser anteaters Tamandua tetradactyla in the cerrado of Brazil. Oryx, 35(2), 166–169. doi:10.1046/j.1365-3008.2001.00162.x

Rytwinski, T., Harper, M., Taylor, J. J., Bennett, J. R., Donaldson, L. A., Smokorowski, K. E., … Cooke, S. J. (2020). What are the effects of flow-regime changes on fish productivity in temperate regions? A systematic map. Environmental Evidence, 9(1), 7. doi:10.1186/s13750-020-00190-z

Sá-Oliveira, J. C., Hawes, J. E., Isaac-Nahum, V. J., & Peres, C. A. (2015). Upstream and downstream responses of fish assemblages to an eastern Amazonian hydroelectric dam. Freshwater Biology, 60(10), 2037–2050. doi:https://doi.org/10.1111/fwb.12628

Sá-Oliveira, J. C., Isaac, V. J., Araújo, A. S., & Ferrari, S. F. (2016). Factors Structuring the Fish Community in the Area of the Coaracy Nunes Hydroelectric Reservoir in Amapá, Northern Brazil. Tropical Conservation Science, 9(1), 16–33. doi:https://doi.org/10.1177/194008291600900103

Salafsky, N., Boshoven, J., Burivalova, Z., Dubois, N. S., Gomez, A., Johnson, A., … Wordley, C. F. R. (2019). Defining and using evidence in conservation practice. Conservation Science and Practice, 1(5), e27. doi:https://doi.org/10.1111/csp2.27

Schneider, M., Biedzicki de Marques, A. A., & Peres, C. A. (2021). Brazil’s Next Deforestation Frontiers. Tropical Conservation Science, 14, 19400829211020472. doi:https://doi.org/10.1177/19400829211020472

Shamseer, L., Moher, D., Clarke, M., Ghersi, D., Liberati, A., Petticrew, M., … Stewart, L. A. (2015). Preferred reporting items for systematic review and meta-analysis protocols (PRISMA-P) 2015: elaboration and explanation. British Medical Journal, 349, g7647. doi:https://doi.org/10.1136/bmj.g7647

SIGEL. (2021). Sistema de Informações Georreferenciadas do Setor Elétrico. Available from Agência Nacional de Energia Elétrica Sistema de Informações Georreferenciadas do Setor Elétrico Retrieved 30 March 2021, from ANEEL https://sigel.aneel.gov.br/Down/

Silva dos Santos, E. (2017). Alterações geomorfológicas no baixo rio Araguari e seus impactos na hidrodinâmica e na qualidade da água. (PhD), Universidade Federal do Amapa. Retrieved from https://www2.unifap.br/ppgbio/files/2018/03/Santos-2017-Tese-de-Doutorado.pdf

Silva, J. M. C. d., Rylands, A. B., & Gustavo, A. B. d. F. (2005). The Fate of the Amazonian Areas of Endemism. Conservation Biology, 19(3), 689–694. doi:https://doi.org/10.1111/j.1523-1739.2005.00705.x

Simões, P. I., Stow, A., Hödl, W., Amézquita, A., Farias, I. P., & Lima, A. P. (2014). The Value of Including Intraspecific Measures of Biodiversity in Environmental impact Surveys is Highlighted by the Amazonian Brilliant-Thighed Frog (Allobates Femoralis). Tropical Conservation Science, 7(4), 811–828. doi:https://doi.org/10.1177/194008291400700416

Stickler, C. M., Coe, M. T., Costa, M. H., Nepstad, D. C., McGrath, D. G., Dias, L. C. P., … Soares-Filho, B. S. (2013). Dependence of hydropower energy generation on forests in the Amazon Basin at local and regional scales. Proceedings of the National Academy of Sciences, 110(23), 9601. doi:https://doi.org/10.1073/pnas.1215331110

Turgeon, K., Trottier, G., Turpin, C., Bulle, C., & Margni, M. (2021). Empirical characterization factors to be used in LCA and assessing the effects of hydropower on fish richness. Ecological Indicators, 121, 107047. doi:https://doi.org/10.1016/j.ecolind.2020.107047

Vié, J.-C., Richard-Hansen, C., & Fournier-Chambrillon, C. (2001). Abundance, use of space, and activity patterns of white-faced sakis (Pithecia pithecia) in French Guiana. American Journal of Primatology, 55(4), 203–221. doi:https://doi.org/10.1002/ajp.1055

